# Carbon substrate type shapes spatial self-organization in a multi-species biofilm community

**DOI:** 10.64898/2026.03.06.709745

**Authors:** Deyong Zhu, Anna J. Svagan, Michael Kühl, Mette Burmølle

## Abstract

Spatial organization is a defining feature of multispecies biofilms and critically influences microbial interactions and emergent community properties. However, understanding and manipulating how microbes assemble into spatially structured biofilms remains challenging because most experimental frameworks emphasize species composition and pairwise interactions, while often overlooking the spatial constraints on biofilms imposed by the environment. In this study, we focus on how carbon substrate type, distinguishing between diffusible sugars and polymeric substrates, affects biofilm self-organization in a four-member synthetic bacterial community (SynCom). Across all tested conditions, the SynCom consistently formed more biofilm biomass than any of its subsets, indicating a robust synergistic phenotype. Using chemically defined, 3D-printed hydrogel substrates with consistent physical properties, we varied carbon source composition to identify its impact on biofilm assembly. Microscopic imaging showed that carbon substrate type strongly influenced biofilm self-organization with diffusible simple carbon substrates yielding relatively intermixed communities, whereas polymer-rich carbon substrates promoted a highly structured biofilm organization characterized by the dominance and peripheral localization of polymer-degrading species. Bioinformatic analyses of carbohydrate-active enzymes (CAZymes) annotation and genome-scale metabolic modeling suggested that metabolite exchange networks in the SynCom may drive more complex metabolic interactions beyond the commonly observed degrader-exploiter-scavenger relationship within planktonic microbial communities. Together, our findings demonstrate carbon substrate type as an important ecological determinant of biofilm self-organization, highlighting the need to integrate environmental factors alongside species composition and metabolic potential to fully understand and manipulate natural and engineered multispecies biofilms.

## Introduction

In natural and engineered environments, microorganisms predominantly grow as spatially structured biofilms that drive emergent ecological and functional properties collectively^1–5^. Within these assemblages, microbes exchange metabolites^6^, compete for limited nutrients^7^, and produce extracellular polymeric substances (EPS)^8^ that confer biofilm resilience to harsh environments and stabilize the community^9–11^. Unravelling how such interactions govern biofilm architecture and function remains a central challenge in microbial ecology^12–14^. Traditionally, this challenge has been approached via studies of which species are present in the community and how they interact metabolically or chemically^15–18^. However, biofilms are fundamentally spatial systems, and recent work has highlighted that the spatial organization of species under specific conditions critically influences their interactions and expressed phenotypes^19–22^.

Resources, particularly the chemical nature of carbon substrates, have long been recognized as a major determinant of microbial interactions^23–25^. For instance, carbon sources influence cross-feeding^26^, niche partitioning^27^, and competition-cooperation balances^28^ across microbial communities. Yet, in natural habitats such as soils, plant surfaces, sediments, and decaying organic matter, microbes rarely encounter carbon substrates as homogeneous, freely diffusible solutes, which are typically employed in many biofilm model systems^29–31^. Instead, they interact with carbon resources that vary widely, with chemical and physical constraints not only in identity and concentration but also in molecular size, solubility, and accessibility^32–35^. For biofilm-associated microbes, such resource properties can generate localized gradients and impose spatial constraints that shape where cells grow, how they interact, and which metabolic strategies are favored^9^.

The spatial complexity of resource landscapes can profoundly alter microbial interactions by i) shaping the local availability of substrates and other metabolites for cells, and ii) constraining or enhancing metabolic dependencies^36–38^. Theory and experimental observations both suggest that polymer degradation, diffusion limitation, and heterogeneous environments can reorganize species distribution and interaction networks in biofilms^39–45^. Particularly, biopolymer degradation in combination with mass-transfer limitations of metabolic substrates and products can generate microscale gradients and localized metabolic niches that select for organized communities of degraders, exploiters, and scavengers^46–49^. Despite extensive studies on polymer-degrading and cross-feeding microbial communities, we still lack a systematic understanding of how carbon substrate type and spatial distribution reconfigures the spatial organization and function of biofilms^50^.

Addressing this knowledge gap requires experimental systems in which community composition is controlled, while the substrate type is systematically varied to enable quantitative investigations of how carbon substrate type alters biofilm spatial assembly. Building on methodological developments in tissue engineering, 3D printing technologies have been used to embed microbial cells within printable hydrogels, or so-called “bioinks”, to create cell-laden constructs^51–54^. These methods are now increasingly used to design and construct living materials as functional engineered systems with defined heterogeneity, e.g., controlling initial microbial placement^51–54^. However, such approaches fundamentally print the initial state, not the emergent biofilm architecture, which requires further maturation into living biofilms^55^. In these cases, the term “3D printed biofilms” is often misunderstood due to the fact that once microbes begin growing, metabolizing and secreting EPS, these printed cell constructs quickly proliferate and remodel their own biofilm architecture, diverging from the predefined pattern far more rapidly than in 3D-printed mammalian cell constructs^56^. As a result, most prior work was not designed to assess how microbes behave when nutrients are provided in forms that mimic embedded, slowly degradable biopolymers prevalent in natural organic matter^55^.

A tractable ecological test of biofilm assembly requires a platform, in which the scaffold architecture is held constant in terms of e.g., surface roughness, porosity, hydration, and diffusion constraints^57–59^, while the type of carbon substrate is the variable of interest, e.g., distinguishing between freely diffusible substrates and polymeric substrates embedded in a matrix that require extracellular degradation and limit localized substrate access. Under such conditions, microbial assembly can unfold naturally through dynamic cell-to-environment and cell-to-cell interactions^60–62^, enabling experimental quantification of how resource-imposed spatial constraints shape “who is where” and “how they interact”. This framework offers an opportunity to interrogate microbial spatial ecology at a level of mechanistic control not previously possible.

Based on this framework, we hypothesized that the type of carbon substrates, independent of community composition, impose spatial constraints that drive distinct biofilm architectures and patterns of interspecies association. To test this, we utilized a four-species synthetic bacterial consortium (SynCom), originally isolated from leaf litter^63^, in combination with 3D-printed substrates as artificial leaves to systematically study the influence of varied carbon resources on biofilm assembly (**Fig. 1A**). We used extrusion-based 3D printing to formulate a reproducible hydrogel matrix, into which a carbon source can be embedded in defined formats, to let the SynCom biofilms unfold naturally. We observed synergistic biofilm formation of the SynCom across exposure to different carbon substrates, highlighting a robust phenotype not observed in monocultures or species subsets. Microscopic images demonstrated that carbon substrate type profoundly modulated the spatial biofilm organization on the printed substrates. Diffusible disaccharides yielded intermixed biofilms, whereas polymer-rich substrates induced a structured biofilm architecture centered around a dominant polymer-degrading species. Carbohydrate-Active enZymes (CAZymes) annotation and genome-scale metabolic modeling (GEMs) further revealed potential metabolic exchange networks beyond the classic degrader-exploiter-scavenger relationship in the SynCom biofilm model. Together, our findings suggest that carbon substrate type is a key determinant of emergent biofilm structure. This further highlights that metabolic interactions within multispecies biofilms cannot be understood independently of the spatial constraints imposed by the environment.

**Fig. 1.**
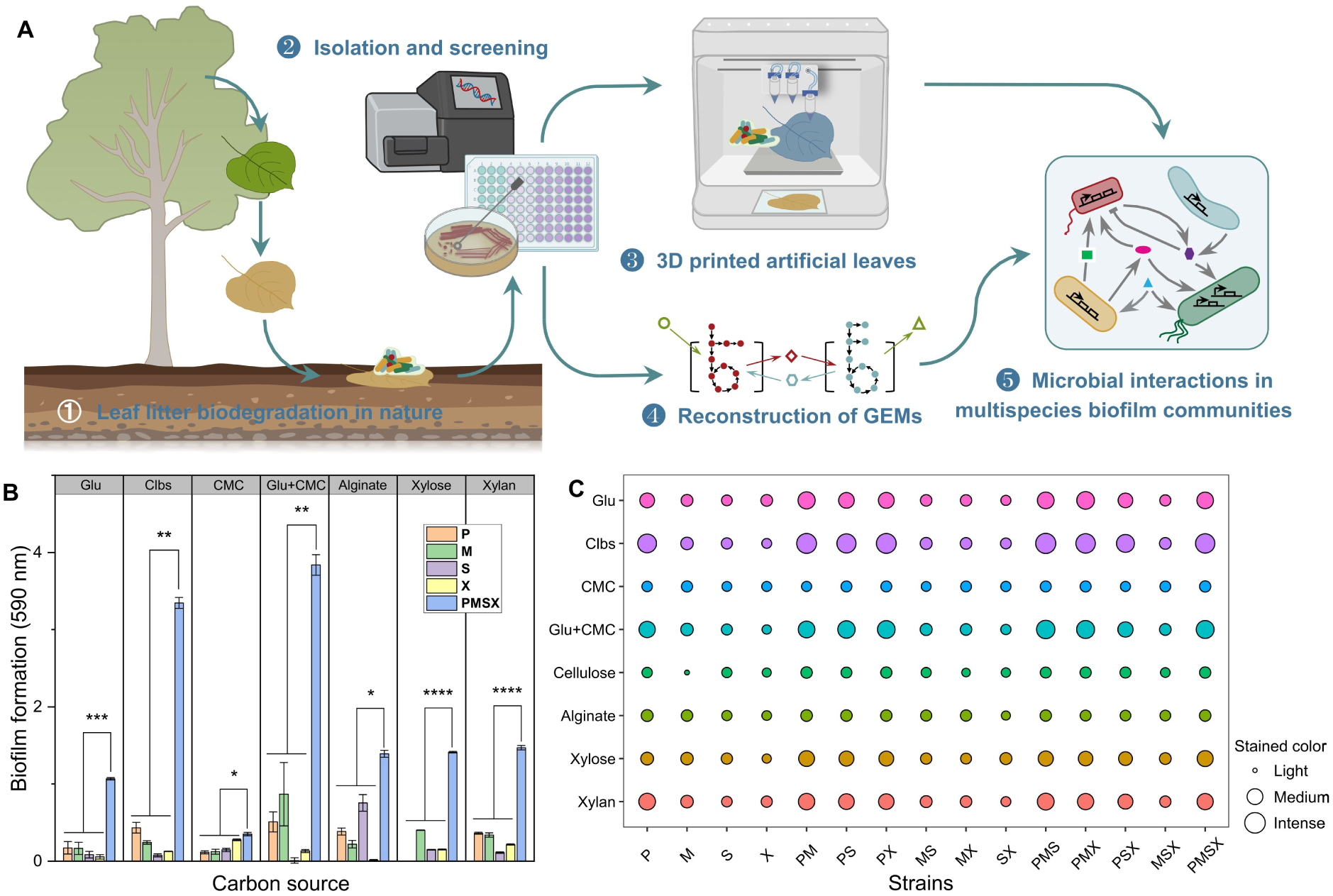
(A) A novel approach using 3D printed artificial leaves and genome-scale metabolic modeling (GEM) for the investigation of microbial interactions and biofilm assembly. (B) Quantification of biofilm formation of monocultures and the SynCom by Crystal Violet (CV) assays in media containing different carbon sources. Abbreviations for carbon sources (Glu: glucose, Clbs: cellobiose, CMC: carboxymethyl cellulose) and strains (P: *P. amylolyticus*, M: *M. oxydans*, S: *S. rhizophila*, X: *S. maltophilia*, PMSX: the SynCom). (C) Screening of the carbon source utilization capacity by mono, double, triple, and quadruple cultures of the SynCom on agar plates. The circle colors represent different carbon sources; and small circles indicate light staining, whereas large ones represent intense staining by Congo red due to biofilm formation on the agar plates.

## Results

### Screening carbon sources for synergistic biofilm formation of the SynCom

We used a four-species SynCom, consisting of *Stenotrophomonas rhizophila* (S), *Stenotrophomonas maltophilia* (X), *Microbacterium oxydans* (M), and *Paenibacillus amylolyticus* (P) in this study (**Table S1**). These bacteria were originally isolated from leaf litter in agricultural soil^63^ and have demonstrated synergistic biofilm formation in rich media such as Tryptic Soy Broth (TSB) in our previous work^64–66^. To assess the influence of different carbon sources on synergistic biofilm formation, the SynCom and monocultures were cultivated in minimal M9 medium supplemented with defined carbon nutrients (**Table S2**), representing common leaf-derived carbohydrate compounds^67^ that range from diffusible sugars (e.g., glucose, cellobiose, xylose) to polymeric long-chain polysaccharides (e.g., cellulose, xylan). Carboxymethyl cellulose (CMC) was tested as an alternative of natural cellulose due to its solubility in aqueous media^68^. Alginate was also included in the screening because it has been widely used in 3D printing as a promising matrix material^69^.

Across all tested carbon sources, the SynCom consistently produced significantly more biofilm biomass than any monocultures, as quantified by Crystal Violet (CV) staining (**Fig. 1B**). The robust phenotype of synergistic biofilm formation under diverse nutritional conditions suggested non-additive interspecies interactions, which cannot be simply extrapolated by the sum of each community member. Among diffusible substrates, cellobiose supported higher total biofilm biomass than glucose or xylose. While CMC alone led to minimal biofilm formation, its combination with glucose substantially enhanced biofilm biomass beyond that observed with only glucose, indicating that polymeric substrates can enhance synergistic biofilm development when combined with readily accessible labile carbon sources. Considering these biofilms were submerged in liquid media, we also tested growth curves, which resulted in altered carbon source utilization capacities under planktonic states compared to biofilms (**Fig. 1B, S1**). For instance, *P. amylolyticus* grew well in both xylose and xylan media, while exhibiting little biofilm formation ability when growing alone. Moreover, we characterized the influence of carbon source concentrations on biofilm formation (**Fig. S2**). The SynCom yielded a remarkable leap in biofilm biomass production from 1 mM to 10 mM for mono- and di-saccharides of glucose, xylose, and cellobiose, while the differences were less obvious for xylan over the range of tested concentrations (0.25 - 1 wt%), and for mono-species (except for *P. amylolyticus* with cellobiose). These data demonstrated that both the presence and concentration of different carbon sources can influence biofilm development.

We used the Congo Red (CR) binding assay to further reveal carbon resource-dependent differences on biofilm formation^70^. Color changes induced by CR binding to biofilm matrix components were recorded by a digital camera followed by digitalized image processing (**Fig. S3**). We observed more intensified staining, indicative of more biofilm formation or altered biofilm matrix composition, with dual-, triple-, and quadruple-species consortia comprising *P. amylolyticus* as compared to mono- or co-cultured consortia without *P. amylolyticus*, indicating a potential dominant role of this bacterium in the SynCom (**Fig. 1C, S3**). Meanwhile, CMC and alginate yielded little color change in all mono- and co-cultures, which was different from the results by CV staining of submerged biofilms (**Fig. 1B, 1C**). A weak color change was noticed with biofilm consortia containing *P. amylolyticus* on the cellulose plates (**Fig. 1C, S3**), while all degradation tests using cellulose filter paper showed negative results with mono-cultures or the SynCom (**Fig. S4**). Together, these results showed that the type of carbon substrate can modulate multispecies biofilm formation, motivating our further investigation into how these variances translate into spatial organization within biofilms under specific conditions.

### Formulation and 3D printing of chemically defined artificial leaves

The screening experiments demonstrated that the SynCom forms biofilm in a non-additive manner across a range of carbon sources; however, these bulk measurements did not capture how the physical type of resources influences spatial organization within multispecies biofilms. To address this limitation, we next developed a scaffold-based system in which architectural features were constant, while different carbon substrates were added in defined physical forms. Because natural leaf litter comprises a heterogeneous mixture of diffusible and polymeric compounds such as simple sugars, cellulose, hemicellulose, lignin, and inorganic minerals^71^, it served as an ecological inspiration for the formulation and 3D printing of chemically defined artificial leaves. To reduce the chemical complexity inherent to natural plant residues while preserving ecological relevance, we fabricated artificial leaf substrates supplemented with defined carbon sources using extrusion-based 3D printing technology (**Fig. 2A**). Cellulose nanofiber (CNF)-based hydrogel was selected as the matrix of printing materials due to its mechanical stability, optical transparency, and compatibility with downstream cell culturing and imaging^72–74^. Importantly, all artificial leaves would share the same CNF backbone, ensuring comparable porosity, diffusion properties, and physical micro-structure across the tested conditions as discussed below.

**Fig. 2.**
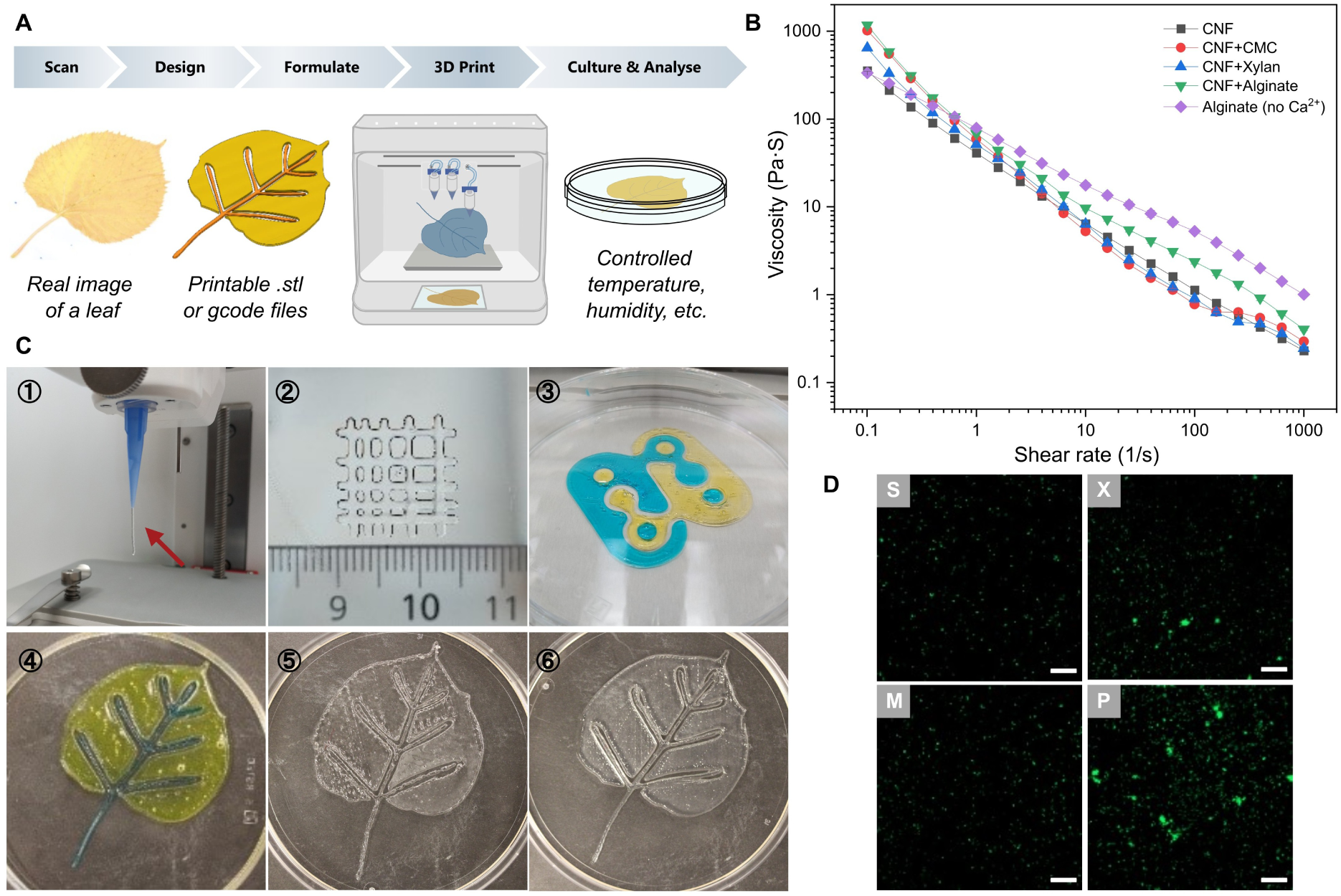
(A) Workflow of 3D printing artificial leaves, including the generation of 3D models by scanning, conversion of the models into printable files, formulation of printable materials, 3D printing, microbial cultivation, and characterization. (B) Rheology property measurements of the formulated hydrogel-based printing materials. CNF: cellulose nanofiber-based hydrogel. CMC: carboxymethyl cellulose. (C) Photographs showing an extruded hydrogel filament from the nozzle (1), a printed grid structure with the smallest square edge of 500 µm (2), a printed dual-color logo of our section (3), a printed dual-color artificial leaf (4), and printed artificial leaves made of varied printing materials of microcrystalline cellulose-based hydrogels (5) and CNF hydrogels (6). (D) Fluorescent images showing the living cells of *S. rhizophila* (S), *S. maltophilia* (X), *M. oxydans* (M), and *P. amylolyticus* (P) by staining with SYTO 9 on the artificial leaf made of the CNF hydrogel alone. Scale bar: 20 µm.

Carbon sources were incorporated into the CNF hydrogel formulation prior to printing (**Table S3**), enabling precise control over chemical composition without imposing spatial patterning of nutrients in the substrates. Rheological characterization confirmed that the addition of diffusible molecules (e.g., glucose, cellobiose) or polysaccharides (e.g., CMC, xylan) caused only minor changes in printability (**Fig. 2B, S5**). Although alginate-Ca^2+^ hydrogels exhibited similar good printability (**Fig. 2B**), we observed precipitation (e.g., Ca_3_(PO_4_)_2_) and degradation of the hydrogel structure when incubated in M9 medium, making the alginate-based hydrogel fragile and not suitable for the subsequent experiments. Atomic force microscopy (AFM) exhibited that the prepared CNF hydrogel comprised 3-4 nm wide and up to 1 µm long nanofibers (**Fig. S6**). Scanning electron microscopy (SEM) revealed highly interconnected porous networks within the CNF-based substrates with pore sizes exceeding typical bacterial dimensions (**Fig. S7**), supporting free diffusion of solutes.

Uniform CNF-hydrogel filaments were extruded using a 3D printer (**Fig. 2C-1**) and the good printability was confirmed by printing a hydrogel-made grid with square edges down to 500 µm (**Fig. 2C-2**). The ability of printing structured shapes was further demonstrated by formulating the CNF hydrogel with different dye solutions and using two extrusion print heads to construct the logo of our research section (**Fig. 2C-3**) and an artificial leaf (**Fig. 2C-4**). Compared to hydrogels made of microcrystalline cellulose (**Fig. 2C-5**), the printed CNF hydrogel yielded smoother edges, more mechanically robust constructs, and better reproducibility of printing (**Fig. 2C-6**). Mono-culture experiments confirmed that the printed CNF-hydrogel substrates were biocompatible, leaving a few island-like living cell aggregates (stained with SYTO 9) on top when only supplying the minimum M9 medium without any carbon sources (**Fig. 2D**). Importantly, no detectable background signal or dye residue was introduced by the CNF-hydrogel substrates during fluorescence microscopy imaging. These properties allowed us to examine how different carbon source formats, rather than the physical structure, shape biofilm assembly on these 3D-printed, reproducible substrates.

**Fig. 3.**
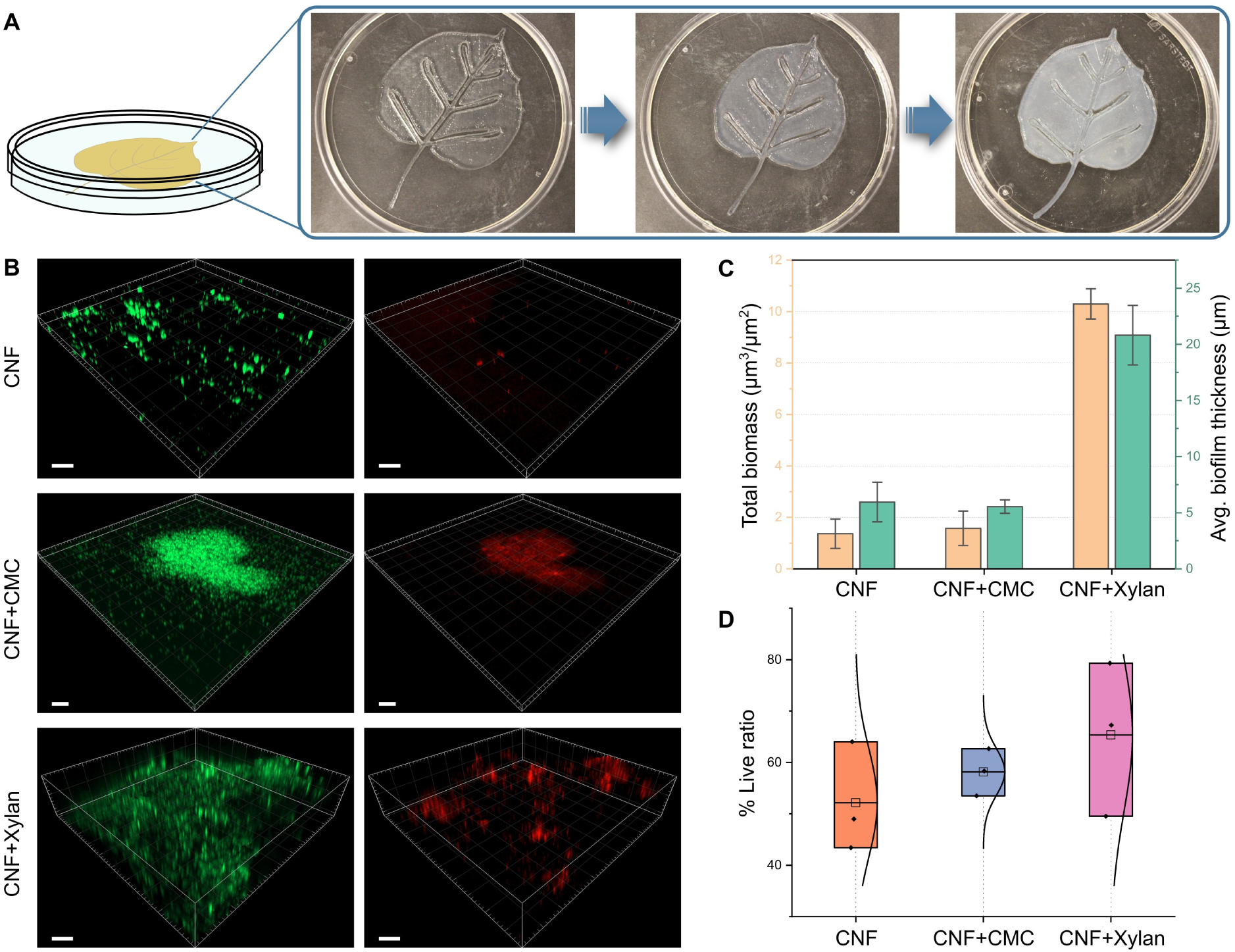
(A) Illustration of the culture system on printed substrates. From left to right: a petri dish container sealed with the gas-permeable, but water-proof membrane and images showing the freshly printed substrate, the substrate after inoculation and with biofilm formed on the top. (B) Fluorescent images showing the live/dead-stained multispecies biofilms formed on the CNF-based substrates containing different carbon sources. CNF+CMC: CNF with carboxymethyl cellulose, CNF+Xylan: CNF with Xylan. Color: green for live cells stained with SYTO 9 and red for dead cells stained with propidium iodide (PI), respectively. Scale bar: 20 µm. (C) Quantification of total biofilm biomass and averaged biofilm thickness. (D) The living cell ratio within each biofilm formed on the substrates with different carbon sources of CNF, CNF+CMC, and CNF+Xylan.

**Fig. 4.**
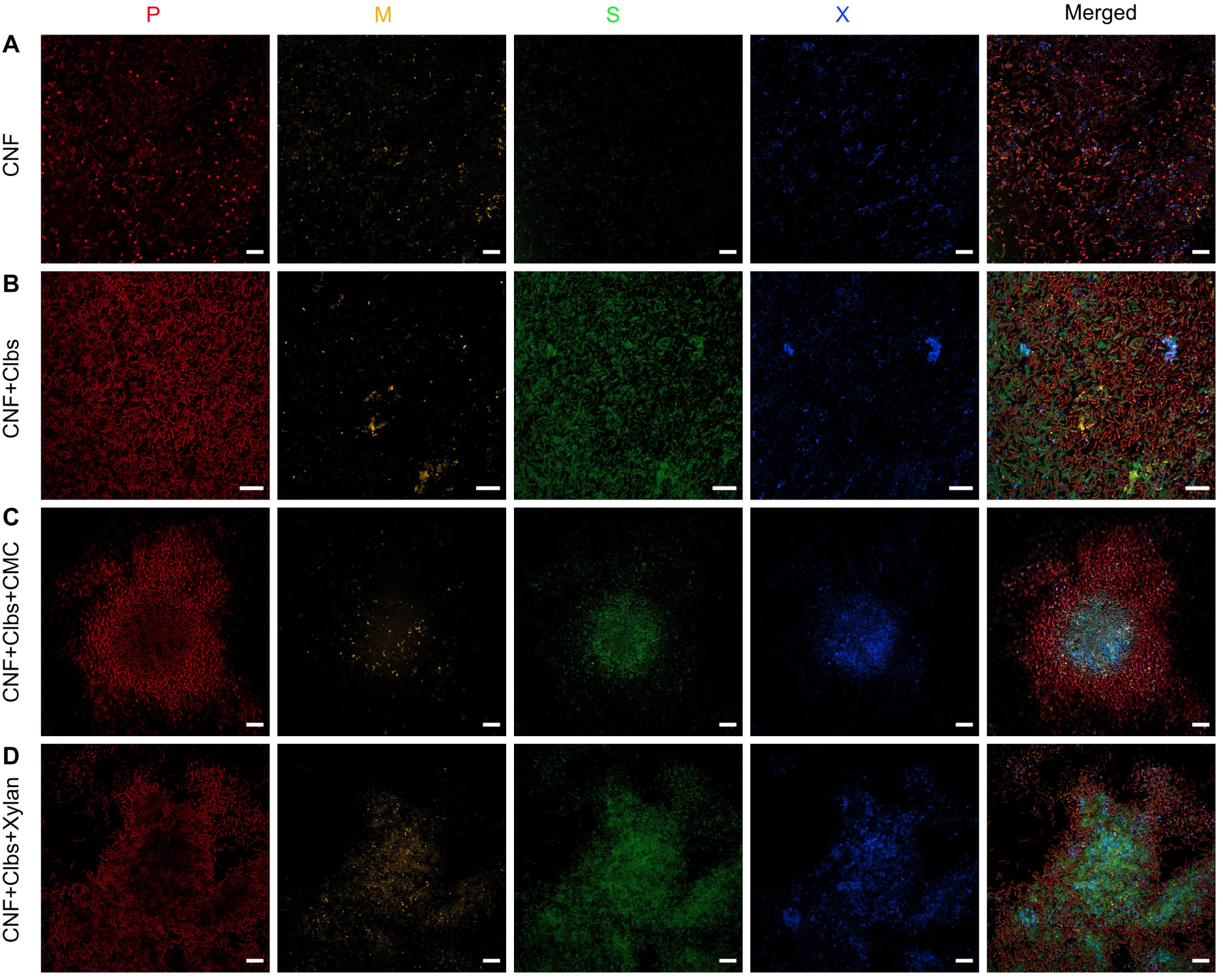
Confocal images showing te spatial structures of the four-species SynCom biofilms on different carbon sources of Panel (A) cellulose nanofiber (CNF), (B) CNF with cellobiose (CNF+Clbs), (C) CNF with Cellobiose and CMC (CNF+Clbs+CMC), and (D) CNF with cellobiose and Xylan (CNF+Clbs+Xylan). Columns with different colors indicate the labeled strains by fluorescence *in situ* hybridization (FISH) with red: *P. amylolyticus* (P), yellow: *M. oxydans* (M), green: *S. rhizophila* (S), and blue: *S. maltophilia* (X). Scale bar: 20 µm.

**Fig. 5.**
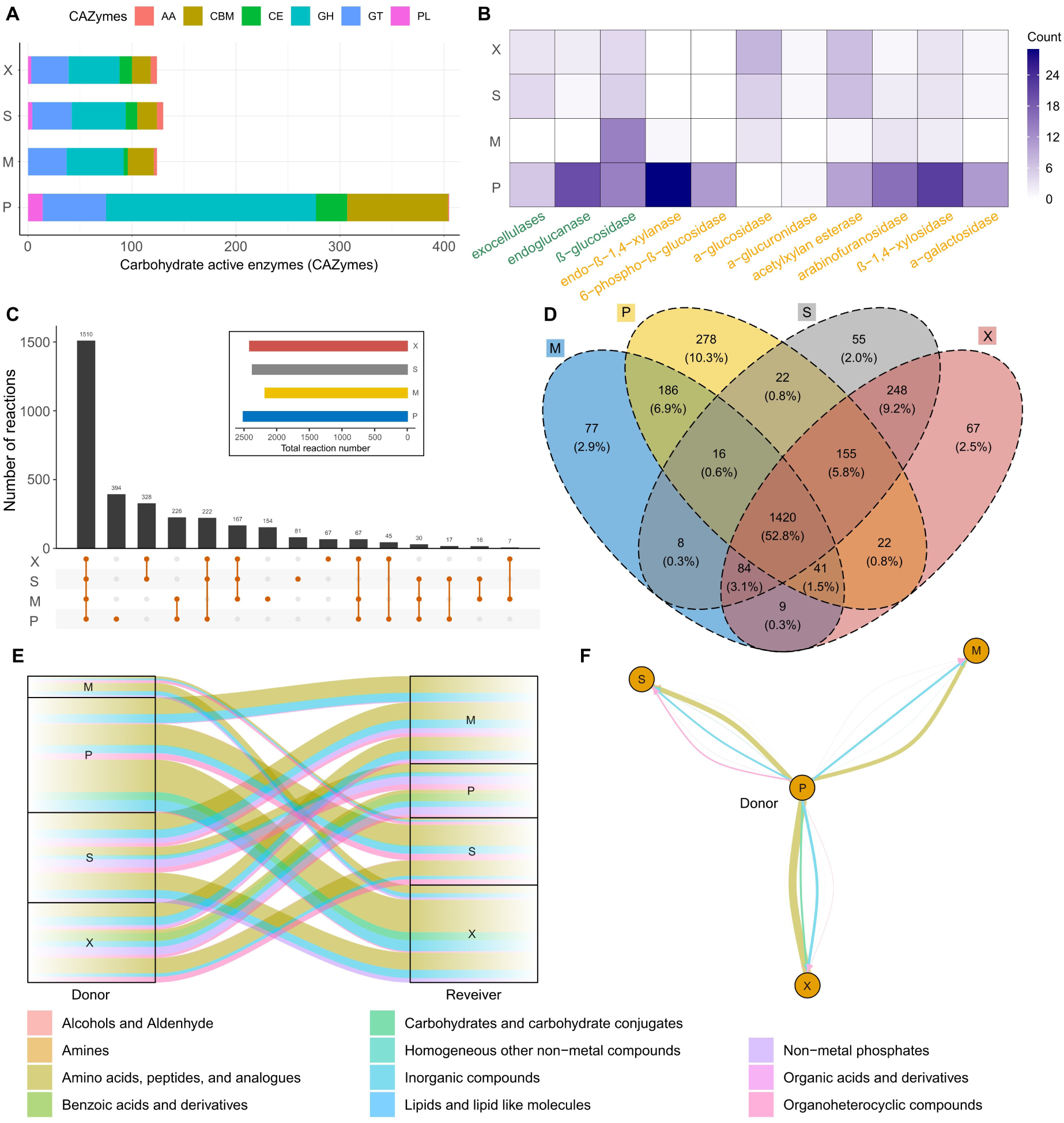
(A) Annotation of Carbohydrate-Active enZymes (CAZymes) including Auxiliary Activity (AA), Glycoside Hydrolase (GH), Glycosyl Transferase (GT), Carbohydrate Esterase (CE), Polysaccharide Lyase (PL) families and Carbohydrate-Binding Modules (CBM) of the four species (P: *P. amylolyticus*, M: *M. oxydans*, S: *S. rhizophila*, X: *S. maltophilia*). (B) Annotated genes that are responsible for degrading cellulose (green) and xylan (orange). (C) The overlap of all reactions predicted by the genome-scale metabolic modeling of the SynCom members. (D) The overlap of all metabolites predicted by the genome-scale metabolic modeling of the SynCom members. (E) Potential metabolic exchanges within the SynCom members simulated by the genome-scale metabolic modeling. (F) Metabolic interaction map centered with *P. amylolyticus* as the donor. The line thickness in (E) and (F) is proportional to magnitude of SMETANA scores and the line color relevant to the classification of compounds as shown at the bottom.

### Carbon source type drives distinct spatial self-organization of multispecies biofilms

To investigate the role of carbon source type on biofilm assembly, instead of predefining the spatial organization of microbial cells embedded in hydrogel matrix as done in earlier studies^51–54^, we incorporated defined carbon sources in CNF hydrogels prior to printing and inoculated microbial cultures on top of the substrates to allow biofilms to grow and mature naturally (**Fig. 3A**). Microscopic imaging by confocal laser scanning microscopy (CLSM) revealed that the spatial organization of SynCom biofilms strongly depended on the carbon source composition (**Fig. 3B**). CNF and CNF+CMC yielded similar total biofilm volume and average biofilm thickness (**Fig. 3C**); however, small and sparse aggregates were formed in the presence of only CNF, while bigger and close-packed biofilms were developed when CMC was present (**Fig. 3B**). Moreover, the addition of xylan supported more extensive and substantially thicker biofilms, as compared to biofilm growth with CNF only (**Fig. 3B, 3C**). Live and dead cell staining results showed comparable overall viability across the tested substrates but indicated uneven distribution of dead cells particularly with xylan (**Fig. 3B, 3D**).

To address the core question of “who is where”, we applied specific oligonucleotide probes labeled with unique fluorochromes (**Table S4**) and used fluorescence *in situ* hybridization (FISH) to enable species-level localization within multispecies biofilms^75^. On CNF-only substrates, biofilm coverage was limited and dominated by *P. amylolyticus* (**Fig. 4A**). The addition of cellobiose, an intermediate during cellulose degradation^76^ that supported the growth of *P. amylolyticus* (**Fig. S1B**), increased total biofilm coverage and maintained the coexistence of multiple species, including *S. rhizophila*, which did not form substantial biofilms on cellobiose alone (**Fig. 1B, 4B**). In contrast, substrates containing both cellobiose and polymeric carbon sources (CMC or xylan) promoted highly structured biofilms, in which *P. amylolyticus* occupied peripheral regions surrounding other community members (**Fig. 4C, 4D**). These observations suggest that the type of carbon substrate strongly regulates spatial organization within multispecies biofilms, where polymeric, less labile substrates favored structured arrangements characterized by localized dominant species.

### Genome-informed prediction of metabolic interaction potential in the SynCom

To further explore the metabolic capacities and potential interactions in the SynCom, we annotated the genomes of each strain against the CAZyme database (dbCAN3)^77^ and revealed substantial differences in carbohydrate-active enzyme repertories among SynCom members (**Fig. 5A**). *P. amylolyticus* encoded an almost three-fold higher number of gene hits covering all six CAZyme families, as compared to the other three species. This includes enzymes implicated in the degradation of cellulose (e.g., endoglucanase (EC 3.2.1.4), exoglucanase (EC 3.2.1.91), β-glucosidase (EC 3.2.1.21)) and xylan (e.g., endo-1,4-β-xylanase (EC 3.2.1.8), acetylxylan esterase (EC 3.2.1.72), arabinofuranosidase (EC 3.2.1.55), β-1,4-xylosidase (EC 3.2.1.37)) (**Fig. 5B)**, and is consistent with *P. amylolyticus* playing an apparent dominant role on polymer-containing substrates^78^. The other three species possessed limited repertoires of CAZymes, indicating that their ability to exploit more complex substrates is limited (**Fig. 5A, 5B**).

We reconstructed genome-scale metabolic models (GEMs) by using the whole-genome sequences of each SynCom member^79^. The predictive models showed that 46.3% of the reactions (**Fig. 5C**) and 52.8% of the metabolites (**Fig. 5D**) overlapped among the four strains composing the SynCom. Notably, the two Gram-positive strains, *P. amylolyticus* and *M. oxydans,* contained more unique metabolic reactions and metabolites related to biochemical pathways involving amino acids and derivatives, carbohydrates, nitrogen and sulfur metabolism, and membrane transportation. On the other hand, more distinct metabolic processes involving fatty acids, lipids, and oxidative stress response were uncovered in the two Gram-negative strains, *S. maltophilia* and *S. rhizophila*. This observation supports the findings from our previous study using the same SynCom, showing accelerated keratin degradation rates and an increased abundance of proteins (e.g., glutathione peroxidase) related to oxidative stress in *S. maltophilia* when co-cultured with other community members^80^. Emergent community-properties may thus arise due to interspecies interactions via activation or deactivation of specific pathways in response to environmental conditions.

GEMs-based analysis by SMETANA further predicted extensive metabolic interactions within the SynCom, with *P. amylolyticus* acting as a major donor of potential metabolic exchanges (**Fig. 5E**). The top-scored metabolites for such inter-species exchange included amino acids, inorganic compounds, carbohydrates, and organic acids. Our analysis indicates the presence of potentially more complex metabolic interaction networks in the SynCom beyond the classic degrader-exploiter-scavenger relationship (**Fig. 5F**). This prediction is consistent with previous metatranscriptomics analyses of the SynCom, showing that differentially expressed genes were highly affiliated to the metabolic processes involving amino acid biosynthesis, membrane transport, and energy conversion^81^.

A high Metabolic Resource Overlap (MRO=0.79, calculated according to Zelezniak *et al.*^82^) among the SynCom members suggested substantial competition for shared resources, particularly in the absence of *P. amylolyticus*, which resulted in an even higher MRO of 0.87. Together, the genomic annotation and metabolic modeling provide a constrained yet informative picture of the metabolic interaction potential within the SynCom. Although these analyses do not directly resolve realized interactions or spatial organization, the predictions suggest that resource-driven metabolic asymmetries may bias how microbial interactions are expressed in space. Elucidating how such metabolic potentials are translated into emergent spatial and functional organization will require future integration of spatially explicit modeling frameworks^4^ and quantitative metabolite measurements^83^.

## Discussion

Despite substantial advances driven by high-throughput sequencing and system-level analyses, general principles governing the assembly of microbial communities particularly in biofilms remain elusive^84^. Biofilms represent the dominant microbial lifestyle in many natural and engineered ecosystems, yet their intrinsic spatial heterogeneity complicates the interpretation of species interactions and community functions^9, 12, 85–87^. In this study, we examined how the type of carbon substrate shapes the assembly, spatial organization, and inferred metabolic interactions of a four-species SynCom. Using 3D-printed CNF-based substrates that mimic aspects of leaf litter degradation, we demonstrated that polymeric carbon sources such as CMC (carboxymethyl group-modified cellulose from plants) and xylan (one type of hemicellulose) profoundly alter biofilm architecture, as compared to biofilm development with more labile and diffusible carbon sources such as cellobiose. While synergistic biofilm formation and viability remained comparable across the tested conditions, the spatial distribution of cells and species differed markedly, indicating an important role of carbon substrate type in modulating the spatial organization of biofilm community members. The type of carbon substrate thus appears as an important ecological variable, which can generate spatial niches in chemically defined and physically consistent environments.

The formation of aggregated and spatially partitioned biofilm structures on polymer-containing substrates likely reflects localized access to degradation products released via extracellular enzymatic activity^9, 11, 88–90^. In contrast to freely diffusible substrates, polymeric carbon sources require enzymatic depolymerization, effectively constraining carbon availability to regions near active degraders^9, 42^. Such constraints can promote spatial segregation and structured biofilm growth, consistent with ecological theory predicting that diffusion limitation and localized resource conversion foster spatial patterning in microbial communities^91–93^. Importantly, such spatial effects arose in our study despite the use of a consistent hydrogel backbone across all printed constructs, indicating that differences in biofilm architecture were driven primarily by chemical resource form rather than physical scaffold heterogeneity. This distinction is critical, as it suggests that even subtle changes in resource chemistry can lead to a spatial reorganization of microbial biofilm communities without altering bulk material properties.

Previous applications of 3D printing in biofilm studies have largely focused on embedding cells or defining their initial spatial arrangements^51–54^. In contrast, our study used 3D printing to standardize the physical environment, while selectively varying chemical resource composition, allowing biofilm structure to emerge naturally through microbial growth and interaction. A limitation of our current artificial leaf design is that it captures only a subset of the chemical and structural complexity of natural leaf litter, instead of recreating leaf surface topography in a microscopically faithful manner^94–96^. While extrusion-based 3D printing offers opportunities to incorporate additional polymers or gradients in future designs, the present system should be viewed as a reductionist, yet tractable platform for dissecting how defined environmental variables influence multispecies biofilms.

Genome annotation and metabolic modeling revealed substantial differences in the CAZymes repertoires among the SynCom members, with *P. amylolyticus* encoding a disproportionately large capacity for polymer degradation. This metabolic potential of *P. amylolyticus* aligned with its dominant spatial positioning in biofilms grown on polymer-containing substrates, where it surrounded or overlaid other species. However, this dominance was not universal across conditions, emphasizing that functional complexity emerges from the interaction between metabolic capacity and environmental context, rather than being an intrinsic property of individual species^93, 97, 98^. Genome-scale metabolic modeling further predicted extensive metabolic overlap among community members, suggesting strong potential for competition. Despite this predicted metabolic overlap, the consortium consistently exhibited synergistic biofilm formation across tested conditions. This suggests that interaction outcomes in structured biofilms cannot be inferred from metabolic network overlap alone, and that spatial organization and localized substrate accessibility may modulate how competitive and potentially synergistic interactions are realized. These findings underscore the importance of considering spatial context when interpreting metabolic interaction networks and community-level phenotypes^99–101^.

However, we also note further improvements of our current framework for metabolic modeling analysis^4, 102–104^. Although the predicted role of *P. amylolyticus* as a major donor of metabolites, coupled with high metabolic resource overlaps, suggests that both cooperation and competition coexist within the community, these predictions remain inherently non-spatial and therefore cannot fully capture how metabolite exchange is realized within structured biofilms. Rather than claiming direct causality between predicted metabolic exchanges and observed spatial patterns, our results support a more nuanced interpretation that carbon substrate type shapes spatial organization, which in turn modulates the expression and consequences of metabolic interaction potential. Resolving how these processes dynamically reinforce one another requires future integration of spatially explicit modeling (e.g., BacArena^105^, COMETS^106^), quantitative metabolite measurements (e.g., stable isotope probing and imaging^107^), and time-resolved analyses (e.g., labeling and spectral imaging^108^).

Overall, our findings suggest that resource-imposed spatial constraints may be a fundamental driver of microbial community organization, cooperation, and function during multispecies biofilm assembly. Our study highlights the necessity of integrating carbon substrate type, spatial organization, and metabolic potential to understand multispecies biofilm ecology. By combining controlled experimental systems with genome-informed modeling, we provide a framework for probing how environmental factors govern microbial community assembly. Further development of spatially resolved experimental and computational tools will be essential for translating these insights into predictive models of biofilm behavior in natural and engineered ecosystems.

## Materials and Methods

### Chemicals and bacterial species

All chemicals were purchased from Sigma-Aldrich unless otherwise noted. The *2,2,6,6-*tetramethylpiperidine-*1*-oxyl (TEMPO)-cellulose nanofiber (CNF)-based hydrogels (TEMPO-CNF, for short as CNF) were prepared as described previously in detail^109^, with the exception that a M-110 EH Microfluidizer processor (Microfluidics Corporation, Westwood, MA, USA) was used. The starting pulp, a never-dried bleached sulfite pulp from spruce (<1% lignin), was a kind gift from Nordic Paper Seffle AB, Sweden.

A consortium of four bacteria, *Stenotrophomonas rhizophila* (S), *Stenotrophomonas maltophilia* (X), *Microbacterium oxydans* (M), and *Paenibacillus amylolyticus* (P), previously co-isolated from leaf litter samples in agricultural soil^8^, was used as the SynCom in this study (**Table S1**). Bacterial isolates from −80 °C preserved stock cultures were streaked on Tryptic Soy Agar (TSA) plates (containing 30 g L^-1^ Tryptic Soy Broth (TSB) and 15 g L^-1^ agar) and incubated for 48 hours at 24 °C until single colonies could be picked. Single colonies were inoculated in 5 mL TSB (30 g L^-1^) in test tubes and incubated overnight at 24 °C with shaking at 250 rpm, which were referred to as overnight cultures and subsequently used in the next steps.

### Biofilm formation quantification and growth curve measurement

#### Crystal Violet (CV) assay

The total biomass of submerged biofilms was measured using a modified CV assay^64^, in which biofilms formed on the pegs of a Nunc-TSP lid system (Thermo Fisher Scientific Nunc A/S, Denmark) were quantified by CV staining. Briefly, overnight cultures of each strain were washed twice by 1x PBS and resuspended in the minimal medium M9 (prepared according to the manufacturer’s instruction and named as M9 in the following) supplemented with different carbon sources as listed in **Table S2**, followed by adjusting OD_600_ to 0.15. CMC was tested as an alternative to cellulose due to its higher water solubility. Additionally, alginate was also tested because it has been reported as a promising matrix material for 3D printing^69^. The synthetic communities were prepared by mixing the cultures at equal volumes. An aliquot of 150 µL of each culture or combination was added per well in a 96-well microtiter plate with three replicates per sample. After 24 h incubation at 24 °C under static condition, the peg lid was transferred sequentially to three washing trays containing 1x PBS to wash off planktonic cells, followed by staining the biofilms formed on the pegs in a 96-well microtiter plate containing 180 µL of 1% (w/v) CV solution per well. After 20 min of staining, the peg lid was rinsed by 1x PBS five times and then placed into a new microtiter plate with 200 µL of 96% ethanol per well. After standing 25 min to allow the bounded CV molecules to dissolve into ethanol, the absorbance of CV in ethanol was measured at 590 nm by the plate reader (ELx808, BioTeck Instrument, VT, USA). Dilution of the CV-ethanol suspension was applied if the absorbance value was above 2.

#### Congo red dye-binding assay

The mono-, double-, triple-, and quadruple-species biofilms formed on agar plates were quantified by the modified Congo red dye-binding assay^70^. The Congo Red agar plates were prepared by mixing 1.5% (w/v) Difco™ Agar Noble (BD, France), 40 µg mL^-1^ Congo red, and M9 supplemented with carbon sources as listed in **Table S2**. Mono-species and combinations of bacterial cultures were prepared as previously described in the CV assay, except that the OD_600_ was adjusted to 0.3. An aliquot of 3 µL from each culture was inoculated on the Congo red-agar plates. All plates were incubated at 24 °C for three days. Images were acquired by a Canon EOS 700D camera and digitally processed by ImageJ (https://imagej.nih.gov/ij/) to quantify the color changes of the colony zone as compared to the background.

### Rheological property measurement

The rheological property of the formulated hydrogel-based 3D printing materials was measured by the Discovery HR-3 hybrid rheometer (Thermal Analysis Instruments, DE, USA) equipped with a 40 mm parallel plate geometry and a measurement gap of 200 µm. Shear rates in the range of 0.1 to 1000 s^-1^ at a frequency of 1 Hz were used to determine the viscoelastic range of the 3D printing materials.

### 3D printing artificial leaves

The formulation and carbon source concentrations of 3D printing materials used in this study are listed in **Table S3**. The printing models of the testing grid and artificial leaves (original images from a lime tree leaf) were generated by Fusion 360 (Autodesk Inc., CA, USA). The printable files were produced and validated using HeartWare (Cellink Inc., Sweden). All models were printed by a commercial 3D bioprinter (BioX G2, Cellink Inc., Sweden) equipped with a 22G/0.41mm nozzle. Printing parameters, including pressure, traveling speed, and infill rate/geometry, were optimized based on the material’s printability. Typical parameters were 15 kPa (pressure), 6 mm s^-1^ (traveling speed), and 95% infilled with honeycomb-like infill geometry for the CNF-based hydrogels. The printed substrates were further post-treated in a laminar flow hood for 4 hr for partial dehydration prior to inoculation.

### Biofilm formation on the printed substrates containing different carbon sources

Mono-species bacterial cultures and the SynCom were prepared as previously described in the Congo red dye-binding assay. The microbial cell suspensions were evenly inoculated onto the artificial leaves containing different carbon sources, followed by incubation in petri dish sealed with gas-permeable but water vapor-proof membranes (Breathe-Easy® sealing membrane, Sigma-Aldrich Labware, Diversified Biotech, MO, USA) under controlled temperature and humidity (24 °C, 60%RH).

### Biofilm characterization by Confocal Laser Scanning Microscopy (CLSM)

#### Live/dead staining

Biofilm samples were washed twice with 0.9% NaCl and then stained by the FilmTracer™ LIVE⁄DEAD Biofilm Viability kit (Invitrogen; Thermo Fisher Scientific, OR, USA) according to the manufacturer’s instructions. After removal of the washing buffer residues, biofilms formed on artificial leaves were *in situ* characterized by CLSM (Zeiss LSM 800 Airyscan) with an LD Plan-Neofluar 40x/0.6 objective using the Quick-LUT function to set the pixel saturation limits. The maximum excitation/emission wavelengths for SYTO 9 and propidium iodide (PI) were 482/500 nm and 490/635 nm, respectively. Images with Z-stacks (layer distance 1 µm) were processed by the free software Imaris Viewer (version 10.0.1, Bitplane AG, Belfast, UK) and Comstat2 (version 2.1, https://www.comstat.dk/)^110^.

#### Fluorescent in situ hybridization (FISH)

A protocol described previously by Liu *et al.*^66^ was applied with some modifications. Briefly, biofilm samples on artificial leaves were fixed in 4% (w/v) paraformaldehyde in 1x PBS overnight at 4 °C. After washing twice in 1x PBS (2 min each), the samples were incubated in the permeabilization solution containing 1 mg mL^-1^ lysozyme at room temperature for 10 min. After another two rounds of washing in 1x PBS (2 min each), the samples were dehydrated by three steps of washing with 50%, 75%, and 100% ethanol, respectively, for 3 min each at room temperature. The hybridization procedure with the target fluorochrome-labeled oligonucleotides probes (**Table S4**) were carried out in a freshly prepared stringency hybridization buffer (0.9 M NaCl, 20mM Tris/HCl (pH 7.2), 9% formamide, 0.05% (w/v) sodium dodecyl sulfate (SDS), and probes at a final concentration of 5 ng each per µl) at 46 °C for 2.5 h. Finally, the samples were washed twice with a prewarmed washing buffer (20 mM Tris/HCl (pH 7.5), 102 mM NaCl, and 0.01% (w/v) SDS) and incubated at 48 °C for 15 min. The samples were ready for imaging by the Zeiss LSM 800 CLSM with a Plan-Apochromat 63x/1.4 oil DIC M27 objective (Excitation/Emission: 402/455 nm (X), 492/518 nm (S), 548/561 nm (M), and 650/673 nm (P)) after rinsing with ddH_2_O.

### Carbohydrate-Active enZymes (CAZymes) annotation

CAZymes domains were identified by dbCAN3^77^ based on the reference genomes available online (**Table S1**). Only gene hits that were predicted in at least two of the three tools were selected for the downstream analysis. E value cut-offs for CAZyme categories were optimized on the basis of results matching with the CAZyme database^77^.

### Genome-scale metabolic modeling

Metabolic models of the four species used in this study were reconstructed and auto-curated using the gapseq pipeline with default parameters^111^. Manual curation was performed by removing duplicate reactions and metabolites with different IDs, filling network gaps, and adding missing metabolites and reactions. Model quality was assessed by the MEMOTE test suite (version 0.13.0)^112^. The likely exchanged metabolites among the community members and metabolic resource overlaps were evaluated using SMETANA (Species METabolic interaction ANAlysis)^82^.

## Data and materials availability

Data and R scripts can be obtained from the authors upon reasonable requests.

## Acknowledgements

We thank Prof. Peter Ulvskov (Department of Plant and Environmental Sciences, University of Copenhagen) for his generous help in the rheology measurements, Dr Nan Yang for sharing experience in the FISH experiments, Mads Frederik Hansen and Dr Henriette Lyng Røder for joining group discussions, and technicians Anette Løth and Ayoe Lüchau for their assistance in the laboratory. This study was supported by the grants from the Villum Foundation (Bioprint: project #35906) and the European Research Council under the European Horizon 2020 research and innovation programme (grant No. 101002208, BioMatrix) to M.B.; and grants from the Independent Research Fund Denmark (grant No. DFF-8022-00301B and DFF-8021-00308B) and the Gordon and Betty Moore foundation (grant No. GBMF9206) to M.K.

## Author contributions

All authors contributed to the study design. D.Z. conducted experiments, performed analyses, created visualizations, and wrote the original manuscript. A.J.S. produced and characterized the TEMPO-CNF hydrogel. M.B. and M.K. supervised the project and acquired funding. All authors reviewed and edited the manuscript.

## Supplementary Materials

Supplementary information, tables, and figures – pdf file

