## Supplementary information for "Carbon substrate type shapes spatial self-organization in a multi-species biofilm community"

Supplementary materials for this article contain the following tables and figures:

**Supplementary Methods**
**Table S1**: List of isolated soil strains in the four-species biofilm consortium (SynCom).
**Table S2**: List of carbon sources for microbial utilization screening.
**Table S3**: List of hydrogel-based printing materials.
**Table S4**: List of fluorescence-labeled oligonucleotide probes for the experiment of fluorescent *in situ* hybridization (FISH).
**Fig. S1**. Growth curve measurement.
**Fig. S2**. Carbon source utilization of different concentrations.
**Fig. S3**. Carbon source screening by the agar plate assay.
**Fig. S4**. Cellulose degradation test with cellulose filter paper.
**Fig. S5**. Rheological property measurement.
**Fig. S6**. AFM image of the cellulose nanofibers (TEMPO-CNFs).
**Fig. S7**. SEM images of the surfaces of 3D printed artificial leaves.

### Supplementary Methods

#### *Growth curve measurement*

Growth curves of the mono- and quadruple-cultures in M9 supplemented with different carbon sources (**Table S2**) were measured using a plate reader (Synergy H1, BioTek Instrument, VT, USA) at 24 °C. 96-well microtiter plates containing bacterial cultures were prepared as described previously in the CV assay. Each sample contained three technical replicates.

#### *Cellulose degradation test on filter paper*

The examination of pure cellulose degradation was conducted using porous filter paper (Whatman paper, Z695068). An aliquot of 300 µL of monocultures and the SynCom culture (with $OD_{600}$ adjusted to 0.3) was inoculated on the autoclaved filter paper strips in each tube, individually. After 6 hrs to allow the bacterial cells to sediment onto the paper's surface, 5 mL minimum media M9 was added to each tube, and all tubes were incubated at 24 °C with continuous monitoring for up to 14 days. Blank filter paper strips with no inoculation were used as control. Each sample contained three technical replicates.

#### *Atomic Force Microscopy (AFM) characterization*

The AFM micrograph of TEMPO-CNF was obtained using a MultiMode 8 (Bruker, Santa Barbara, CA, USA) in PeakForce QNM mode using ScanAsyst cantilevers (Bruker, Camarillo, CA, USA). The silicon wafers were first treated by adsorbing a layer of Polyethylenimine (PEI, 0.01 wt%, 1 min), followed by rinsing with Milli-Q water and drying. A drop of the TEMPO-CNF suspension (0.01 wt%) was deposited on the wafer, followed by immediate rinsing by MilliQ-water and drying.

#### *Scanning Electron Microscopy (SEM) characterization*

The morphology of 3D-printed substrates comprising different carbon sources was characterized by SEM. Briefly, the hydrogel-based substrates were pretreated by immersion into liquid nitrogen until completely frozen, followed by lyophilization to preserve the structure. The lyophilized samples were sputter-coated with gold and multiple micrographs were obtained at selected regions using a Quanta 200 environmental scanning electron microscope (FEI, OR, USA). Images at 300x magnification, 5 kV were presented to demonstrate the microscale porous structure.

**Table S1**: List of isolated soil strains in the four-species biofilm consortium (SynCom).

| Strain | Abbreviation | Gram | Reference genome in ENA (European Nucleotide Archive) |
|---|---|---|---|
| *Stenotrophomonas rhizophila* | S | Negative | PRJEB15263 |
| *Stenotrophomonas maltophilia** | X | Negative | PRJEB18431 |
| *Microbacterium oxydans* | M | Positive | PRJEB15265 |
| *Paenibacillus amylolyticus* | P | Positive | PRJEB15262 |

*Genome-wide taxonomy analysis by Genome BLAST Distance Phylogeny (GBDP) shows this genome should refer to *Stenotrophomonas maltophilia* instead of *Xanthomonas retroflexus* in ENA.

**Table S2**: List of carbon sources for microbial utilization screening

| Carbon source name | Abbrev. | Final Conc. | Carbon source type | CAS-no. | Purchased from |
|---|---|---|---|---|---|
| D-(+)-Glucose | Glu | 10 mM | Monosaccharide | 50-99-7 | Sigma-Aldrich |
| D-(+)-Xylose | Xylose | 12 mM | Monosaccharide | 58-86-6 | Sigma-Aldrich |
| D-(+)-Cellobiose | Clbs | 5 mM | Disaccharide | 528-50-7 | Sigma-Aldrich |
| Carboxymethyl cellulose | CMC | 1.0 wt% | Polysaccharide (cellulose) | 9004-32-4 | Sigma-Aldrich |
| Microcrystalline cellulose | Cellulose | 1.0 wt% | Polysaccharide (cellulose) | 9004-34-6 | Sigma-Aldrich |
| Sodium alginate | Alginate | 1.0 wt% | Polysaccharide (alginate) | 9005-38-3 | Sigma-Aldrich |
| Xylan (from corncob) | Xylan | 1.0 wt% | Polysaccharide (hemicellulose) | 9014-63-5 | Carl Roth |

**Table S3**: List of hydrogel-based printing materials

| Material name | Hydrogel composition | Manufacturer of materials |
|---|---|---|
| CNF | 1 % cellulose nanofiber | Home-made for all CNF below |
| CNF+CMC | 1 % cellulose nanofiber mixed with 1 % CMC | Sigma-Aldrich |
| CNF+Xylan | 1 % cellulose nanofiber mixed with 1 % Xylan | Carl Roth |
| CNF+Alginate | 1 % cellulose nanofiber mixed with 1 % sodium alginic acid | Sigma-Aldrich |
| Alginate (no Ca2+) | 1 % sodium alginic acid | Sigma-Aldrich |
| CNF+CMC+Glu | 1 % cellulose nanofiber mixed with 1 % CMC and 10 mM Glucose | Sigma-Aldrich |
| CNF+CMC+Clbs | 1 % cellulose nanofiber mixed with 1 % CMC and 5 mM Cellobiose | Sigma-Aldrich |
| Native cellulose | 5 % native cellulose crosslinked by epichlorohydrin (1 ml in 25 ml) | Sigma-Aldrich |
| CNF+Clbs | 1 % cellulose nanofiber mixed with 5 mM Cellobiose | Sigma-Aldrich |
| CNF+Xylan+Clbs | 1 % cellulose nanofiber mixed with 1 % Xylan and 5 mM Cellobiose | Sigma-Aldrich |

**Table S4**: List of 16S rRNA-directed fluorescence-labelled oligonucleotide probes for the FISH experiment

| Probe | Target | Sequence of probe (5' -> 3') | Fluorochrome |
|---|---|---|---|
| S-Fam | *S. rhizophila* | CGG GTA TTA GCC GAC TGC TT | FAM |
| X-Blu | *S. maltophilia* | CCG TCA TCC CAA CCA GGT ATT | Pacific Blue |
| M-Cy3 | *M. oxydans* | CAT GCG TGA AGC CCA AGA C | Cy3 |
| P-Cy5 | *P. amylolyticus* | CGG TCA GAG GGA TGT CAA GAC | Cy5 |

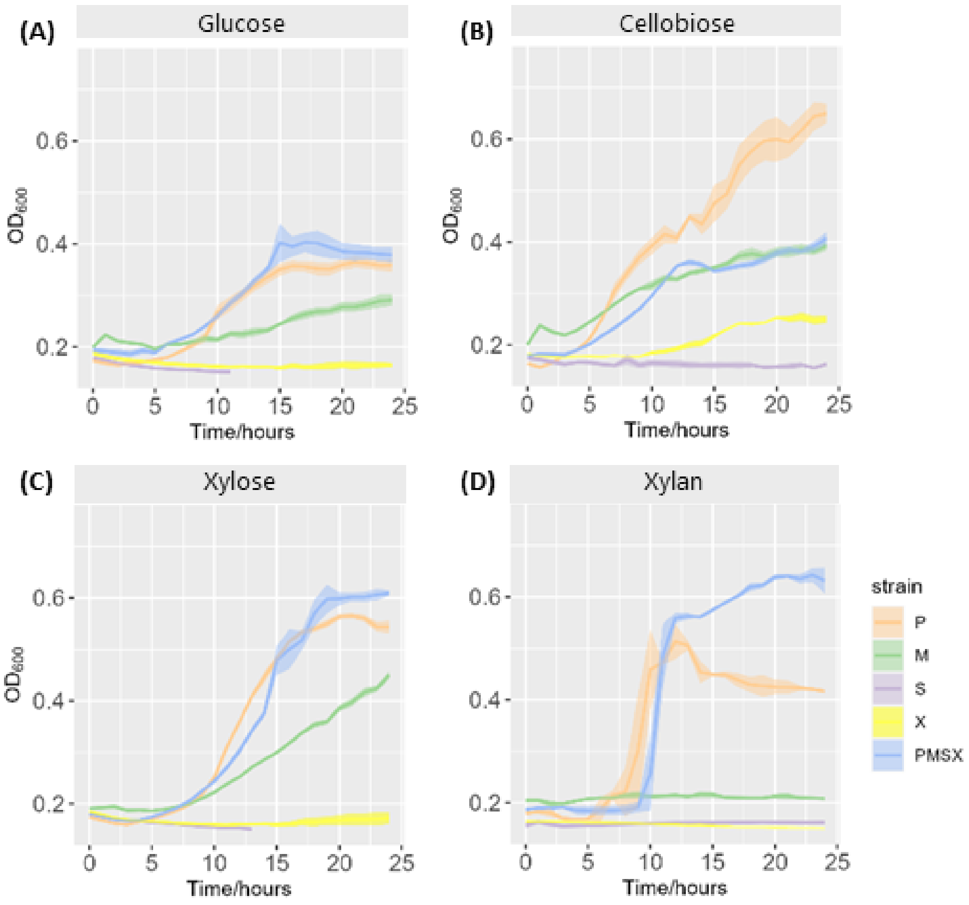


**Fig. S1**. Growth curve measurement in the M9 medium with different carbon sources. (A) Cellobiose, (B) Glucose, (C) Xylose, and (D) Xylan. Each plot represents three technical replicates.

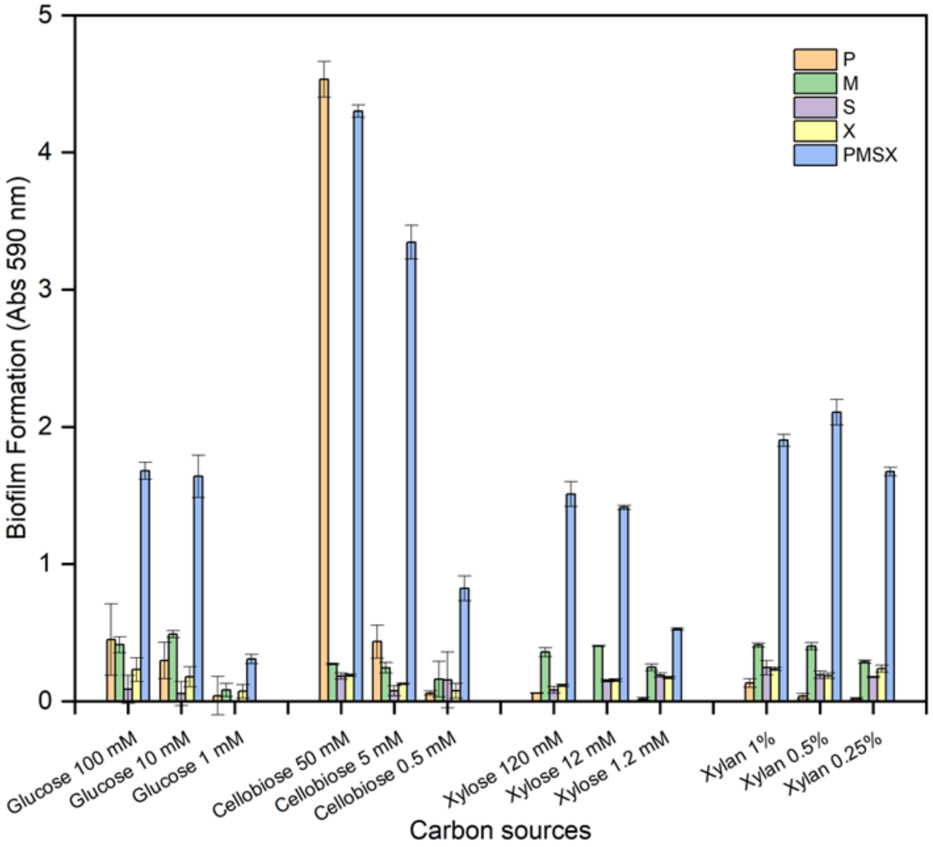


**Fig. S2**. Screening of different-concentration carbon sources by the crystal violet (CV) assay for mono- and multi-species cocultured biofilm formation. S: *S. rhizophila*, X: *S. maltophilia*, M: *M. oxydans*, and P: *P. amylolyticus*.

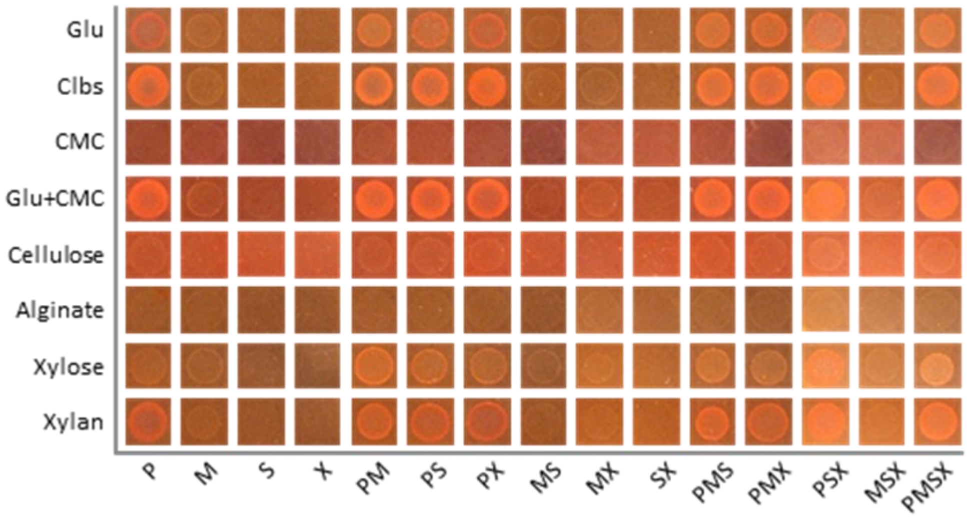


**Fig. S3**. The screening of carbon sources on agar plates with different carbon sources for mono-, dual-, triple-, and quad-species biofilm formation. Glu: glucose; Clbs: cellobiose; CMC: carboxymethyl cellulose; Glu+CMC: glucose with carboxymethyl cellulose.

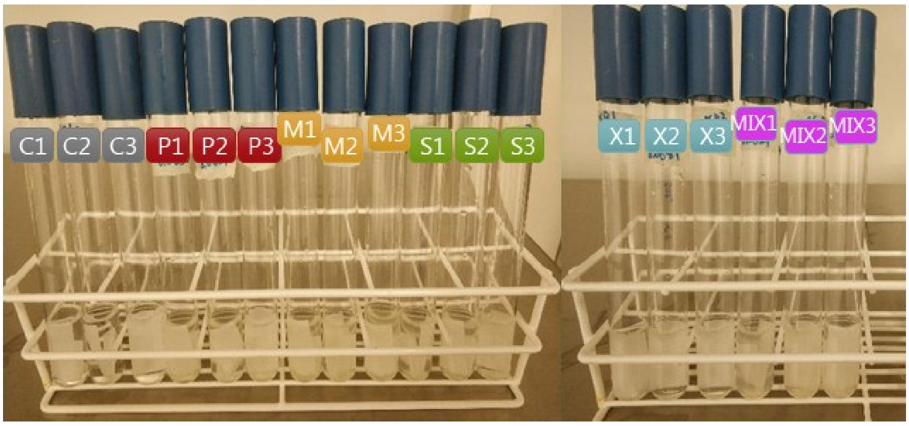


**Fig. S4**. Images of cellulose degradation test with porous filter paper. Continuous monitoring up to 14 days revealed no observable change either in the media or on the filter paper. C: Control without inoculating any bacteria, S: *S. rhizophila*, X: *S. maltophilia*, M: *M. oxydans*, P: *P. amylolyticus*, and MIX: the full consortium. Three replicates of each.

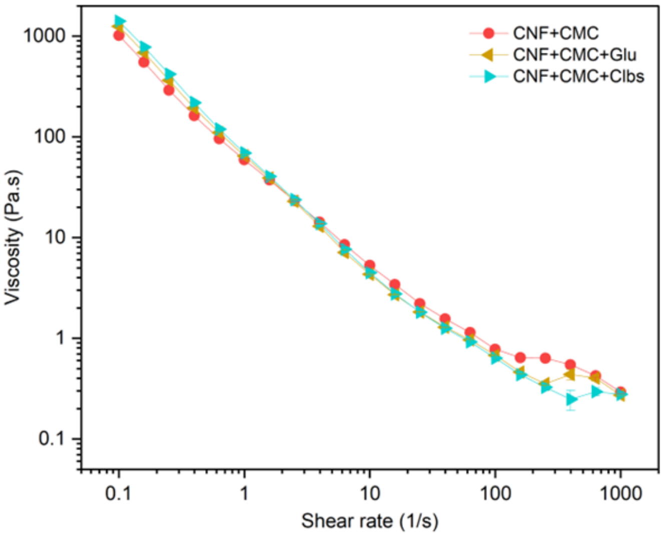


**Fig. S5**. Rheological property measurement of 3D printing materials. CNF: *2,2,6,6*-tetramethylpiperidine-*1*-oxyl-cellulose nanofiber; CMC: carboxymethyl cellulose; Glu: glucose; Clbs: cellobiose. Each data point represents three technical replicates.

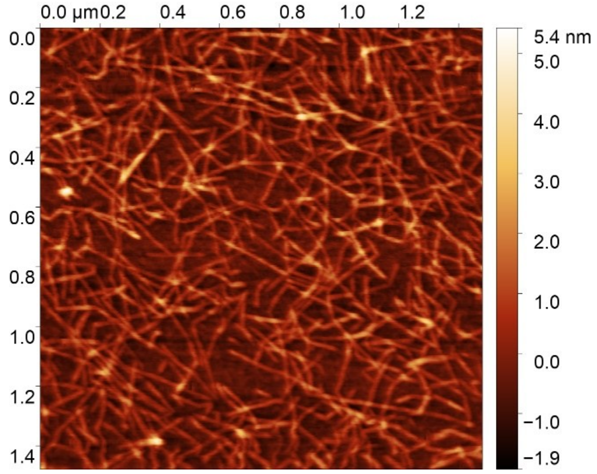


**Fig. S6**. The image by atomic force microscopy (AFM) showing the size of the prepared CNF (TEMPO-cellulose nanofiber).

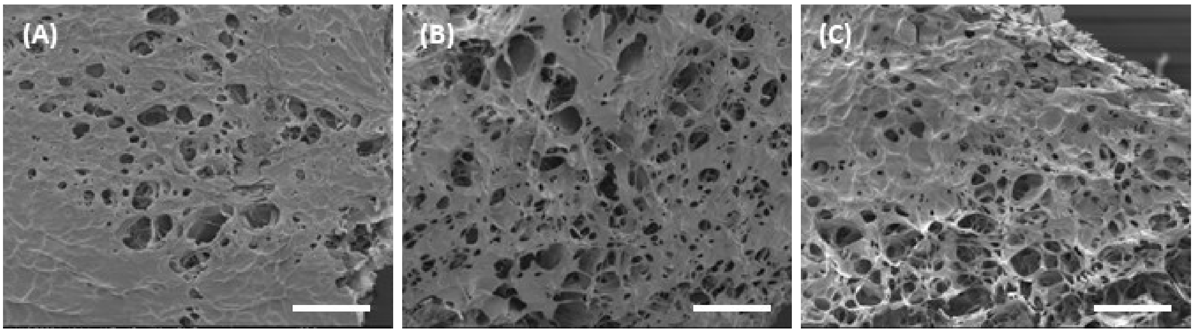


**Fig. S7**. Images by scanning electron microscopy (SEM) demonstrating the interconnected porous properties of printed artificial leaves with carbon sources of (A) CNF, (B) CNF+CMC, and (C) CNF+Xylan. Scale bar: 200 μm. The hydrogels were freeze-dried prior to imaging.
